# Variation in female-biased sexual size dimorphism of Northern Pike (*Esox lucius*) associated with environment and life history

**DOI:** 10.1101/2023.03.06.531313

**Authors:** P.J. Kennedy, M.D. Rennie

## Abstract

**Background:** Sexual size dimorphism (SSD) is a widespread phenomenon in the animal world resulting from differential selection on the sexes. The Northern Pike (*Esox lucius*) is a freshwater apex predatory fish species that exhibits female-biased SSD, but the degree to which SSD varies among populations and what variables might dictate variation in SSD in this species remain poorly understood.

**Aim:** We sought to quantify the degree of variation in SSD among Northern Pike populations across a large portion of their North American range, as well as evaluate associations between the magnitude of SSD in Northern Pike populations with environmental variables and life history traits of populations.

**Methods:** We quantified SSD in 102 populations of Northern Pike across the province of Ontario, Canada, using a standardized gillnetting database. We further investigated the degree to which both environmental variables (Cisco abundance as catch-per-unit-effort, lake surface area, and latitude) and Northern Pike life-history traits (early growth and mortality rates) explained variation in female-biased SSD using linear models.

**Results:** Female-biased SSD in mean weight of Northern Pike increased with increasing Cisco (*Coregonus artedi*) abundance, and the difference in female and male mean age increased with increasing latitude. Furthermore, SSD was greater in populations with lower female mortality and early growth rates.

**Conclusion:** This study indicates that slow-growing, long-lived populations of Northern Pike should exhibit greater female-biased SSD, and that these conditions may be facilitated by the availability of large, energy-dense prey and cooler temperatures at northern latitudes.

## INTRODUCTION

Sexual dimorphism occurs throughout the natural world and can be described as a condition where sexes of the same species exhibit differences in their phenotypic traits in addition to differences in their sex organs (Darwin, 1871). Although sexual dimorphism results in a variety of differences in the phenotypic traits of sexes, a common example is differences in their growth rates and body sizes (Fairbairn *et al*., 2007). This is known as sexual size dimorphism (SSD), and together with other forms of sexual dimorphism, it is often a result of differential selection on the sexes (Fairbairn, 1997; Blanckenhorn, 2005).

While a variety of hypotheses have been proposed for the explanation of SSD, sexual selection through mate choice/competition and fecundity selection are largely considered the driving forces of this phenomenon (Hedrick and Temeles, 1989; Fairbairn, 1997; Blanckenhorn, 2005). When expressed in mammalian and avian species, sexual selection in the form of intra-sexual competition and female choice typically results in males exhibiting larger body sizes than females (Clutton-Brock *et al*., 1977; Cabana *et al*., 1982; Blanckenhorn, 2005). Alternatively, fecundity selection results in SSD biased toward the sex that invests more energy into reproduction through processes such as gonadal development, nest defence, and competition for mates (Parker, 1992). In fishes that express SSD, both male- and female-biased SSD have been observed, and SSD is likely related to breeding systems and ecological differences between the sexes (Roff, 1983; Shine, 1989; Parker, 1992; Pyron, 1996). Male-biased SSD in some fishes appears to be related to male parental care and territoriality over resources or mates (Roff, 1983; Horne *et al*., 2020). Meanwhile, female-biased SSD in fishes is related to strong fecundity selection and may be more likely to occur among species that are more pelagic or bathypelagic and provide no parental care (Horne *et al*., 2020). Further, males and females often exhibit sex-specific habitat segregation in populations with SSD, which can also lead to sex-specific diets and differences in other life-history traits (Ruckstuhl, 2007). Overall, female-biased SSD seems to occur more frequently in fishes (Breder and Rosen, 1966), though the magnitude of SSD observed and the proximate mechanisms driving female-biased SSD have been difficult to discern.

One such fish is the Northern Pike (*Esox lucius*), a widespread apex predator that displays female-biased SSD (Scott and Crossman, 1973; Craig, 1996), though the ecological and evolutionary factors related to variation in SSD observed in this species have yet to be investigated. The growth of male and female Northern Pike has been found to be similar pre-maturation, but females often mature at larger sizes than males, resulting in greater adult female body sizes (Malette and Morgan, 2005). Thus, the magnitude of SSD in this species should be sensitive to adult mortality rate, particularly female mortality rate, because adult mortality is negatively related to body size (Parker, 1992; Charnov *et al*., 2013). Because high adult mortality is correlated with rapid early growth, early maturation, and smaller asymptotic sizes, populations experiencing higher female mortality rates relative to males should have a lower degree of female-biased SSD (Beverton and Holt, 1959; Parker, 1992).

Several aspects of the ecology of Northern Pike suggest that the degree of SSD displayed in this species may also be related to the availability of large-bodied prey. Female Northern Pike have been observed to forage more frequently, display greater post-maturation growth, and invest more energy in gonadal development than males (Diana, 1983; Craig, 1996). Previous work has also indicated that larger Northern Pike occupy deeper waters and have greater versatility in their habitat selection than smaller Northern Pike (Chapman and Mackay, 1984; Pierce and Tomcko, 2005). Considered together, these observations suggest that Northern Pike may exhibit sex-specific habitat segregation, with larger females occupying deeper waters which may be associated with differences in diet and energetics. For instance, larger Northern Pike are known to occupy deeper offshore habitats in lakes with higher densities of Cisco [*Coregonus artedi* (Kennedy *et al*., 2018)]. If larger (putatively female) Northern Pike are better able to exploit large, energy-dense offshore prey (i.e. Cisco) in lakes where they are available, they may experience greater growth efficiency compared to smaller (male) Northern Pike (Pazzia *et al*., 2002). Selection for such behaviour (e.g. utilization of habitats with large-bodied prey) would benefit female fitness explicitly, since body size scales positively with clutch size, and larger numbers of eggs is correlated with greater offspring survival (Roff, 1992; Stearns, 1992).

Our objectives for this study were to first document the degree of variation in SSD among Northern Pike populations across a large portion of their North American range using data from a standardized netting program. Second, we sought to evaluate associations between the magnitude of Northern Pike SSD observed among sampled populations with environmental variables (specifically, Cisco density, lake surface area, and latitude) and life-history traits among populations (mortality and early growth rate indices). We hypothesized that Northern Pike would exhibit greater female-biased SSD in large, high-latitude lakes with greater Cisco densities, as these conditions are predicted to promote longer lifespans and post-maturation growth in this and other apex predatory species (Kaufman *et al*., 2009; Kennedy *et al*., 2018; Lester *et al*., 2021). Additionally, we hypothesized that female-biased SSD would be greater in Northern Pike populations exhibiting lower female mortality and early growth rates. We expected SSD estimates to be more strongly influenced by female life-history traits as females are the sex that typically exhibits greater body sizes, and for whom body size is directly related to fitness via fecundity/egg production (Parker, 1992).

## METHODS

### Data collection and preparation

Data from a standardized gillnetting program, the Fall Walleye Index Netting (FWIN) survey, was obtained from the Ontario Ministry of Natural Resources for our analyses of Northern Pike SSD and prey fish densities. The FWIN dataset included data on inland Ontario lakes that were sampled from 1993 to 2003 using a stratified random sampling design that consisted of overnight sets of multi-mesh gillnets (Morgan, 2002). Sampling occurred in the fall when surface water temperatures were 15°C or cooler and continued until temperatures fell below 10°C. Gillnets were 61 m long, had eight sequential mesh panels (25, 38, 51, 64, 76, 102, 127, and 152 mm) that were each 7.6 m long, and were set perpendicular to the shore for 24 hours at two different depth strata: 2–5 m and 5–10 m. The location and number of nets set were based off the surface area and depth of the lakes with a minimum of eight nets set for each lake. Fish captured were assigned a unique identification number, and the net number, mesh size, location, and date were recorded. Species, fork length, total length, round weight, sex, and age were recorded for all Northern Pike caught. Cleithra and scales were used to estimate the age of Northern Pike. Additional details on the FWIN protocol can be found in Morgan (2002).

### Estimation of SSD response variables

We calculated the mean fork length-at-age of Northern Pike for lakes with at least three individuals in an age class for each sex to compare the average size-at-age of females and males across Ontario. Additionally, we calculated the asymptotic fork length (*L*_10_, mm), mean weight (g), and mean age (years) of female and male Northern Pike from waterbodies with at least 20 captured individuals of each sex to use for response variables. Estimates of *L*_10_ for female and male Northern Pike in each population were calculated as the mean fork length of the largest 10% of the catch for each sex in a lake. This yielded 102 populations with estimations of *L*_10_, mean weight, and mean age (Fig. 1, Table A1). Despite these measures being correlated (Table A2), we included all these response variables for robustness of analyses given possible differences in body shape, growth rates, and body condition among populations that might result in subtle but important differences among relationships with response variables investigated.

**Fig. 1.**
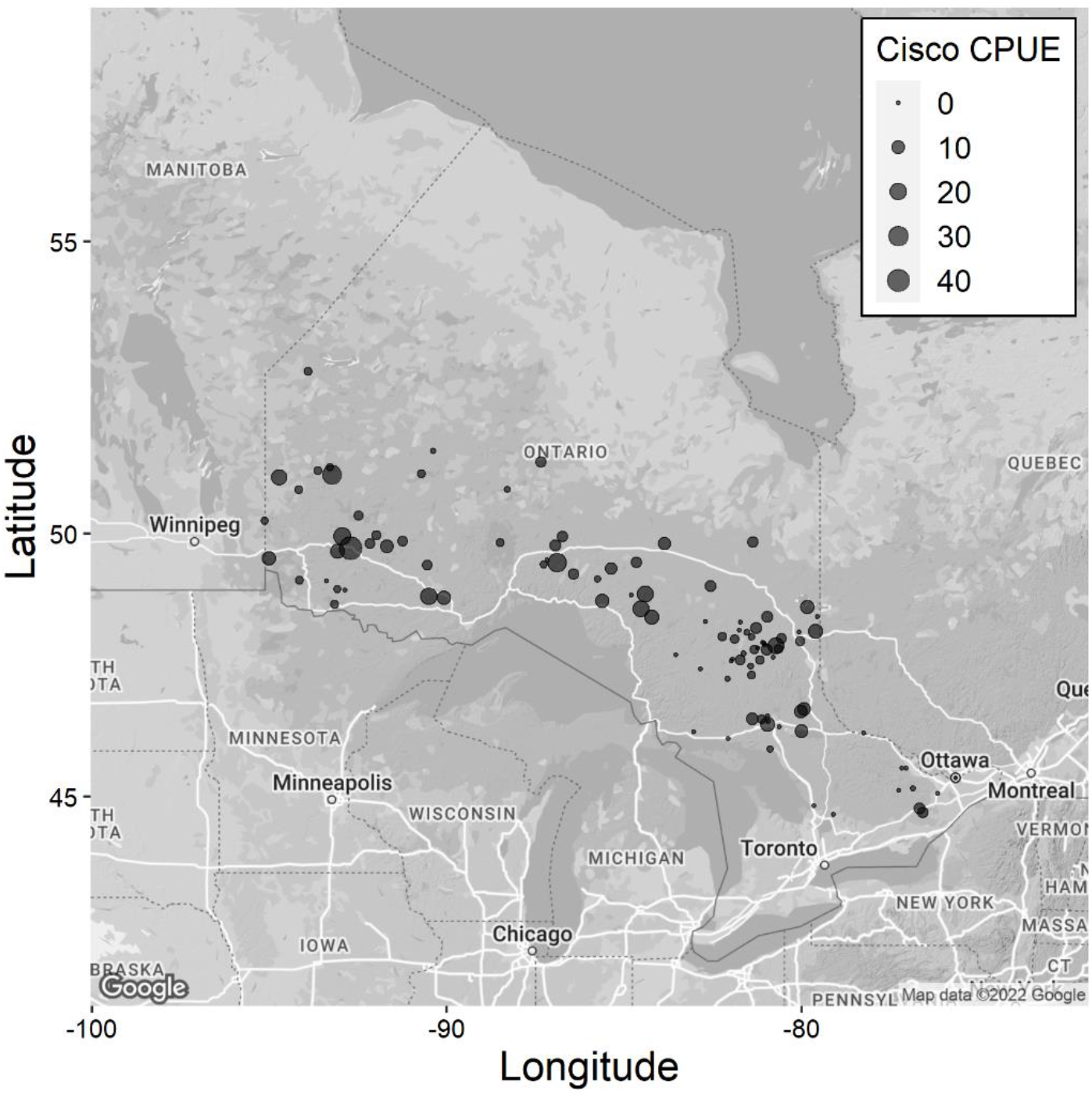
Map of Northern Pike lakes (*n* = 102) by Cisco CPUE. Larger points have greater Cisco CPUE.

Dimorphism for *L*_10_, mean weight, and mean age was calculated using the size dimorphism index (SDI), which is defined as the ratio of the larger (female) to smaller (male) sex minus 1 (Lovich and Gibbons, 1992), which conveniently sets the neutral value (i.e., no dimorphism) to zero. This method presents an intuitive way of quantifying sexual dimorphism, as a value of 0.25 would indicate that females are 25% larger (or heavier, older) than males for the variable of interest. The SDI for mean age of Northern Pike was included to determine if longevity differs between the sexes and whether this is driven by environmental or life-history variables.

### Estimation of environmental predictor variables

Environmental predictor variables included in SSD analyses were Cisco catch-per-unit-effort (CPUE), lake surface area, and latitude. Cisco CPUE was included as a measure of high-quality prey density for adult Northern Pike due to their relatively large size and energy density (Bryan *et al*., 1996), and Northern Pike asymptotic size has been shown to be positively related to Cisco CPUE (Kennedy *et al*., 2018). Lake surface area was used as a measure of overall habitat size, and latitude was used to represent a gradient in both climate and land-use. Cisco CPUE was calculated as a lake-wide area-weighted CPUE, where CPUE and benthic area were calculated in each sampled depth stratum; CPUE was then estimated as an area-weighted lake-wide mean across all sampled strata. For lakes sampled in multiple years, CPUE values were averaged. Lake surface area data were compiled from the OMNRF Aquatic Habitat Inventory (Dodge *et al*., 1985).

### Estimation of life-history predictor variables

Life-history traits included as predictor variables in SSD analyses were female Northern Pike instantaneous total mortality rate (*Z*) and the Gallucci-Quinn Index of early growth rate (ω). Female *Z* was estimated using the Chapman-Robson method (Robson and Chapman, 1961; see also Ricker, 1975), which has been shown to perform better than other commonly employed catch-curve methods for estimating total mortality (Smith *et al*., 2012). First, annual adult survival (*S*) was calculated as:

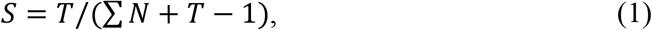

where ∑*N* (∑*N = N*_*x*_ *+ N*_*x* + 1_ *+ N*_*x* + 2_ *+* …) is the total number of fish on the descending limb of the catch curve whose age is equal to the modal age class plus one year (*N*_*x*_). *T* is the sum of the products between recoded age classes and the number of fish caught in the associated recoded age classes, where ages are recoded such that *N*_*x*_ is 0 and the subsequent age class (*N*_*x* + 1_) is 1, and so on. Instantaneous total mortality rates for populations were then calculated as:

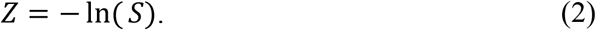

A visual representation of the descending limb of a catch curve (Fig. A1) and an example calculation of *S* and *T* estimates (Table A3) are provided in the Appendix (evolutionary-ecology.com/data/3224Appendix.pdf). Female Northern Pike ω was estimated to represent growth rates near time zero at the size of maximum growth, and was calculated as the product of *L*_*∞*_ (mm) and *K* (per year) (Brody, 1945; Gallucci and Quinn, 1979; Charnov, 2010), which were derived from the von Bertalanffy growth model fit to fork length-at-age data (Ricker, 1975):

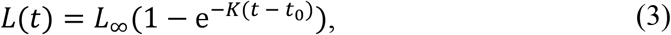

where *L*(*t*) is the fork length (mm) at age *t* (years), *L*_*∞*_ is the asymptotic fork length, *K* is Brody’s growth coefficient, and *t*_0_ is the theoretical age when the fish would have been zero length (assumed to be 0 in our case). In this dataset, female Northern Pike ω was positively correlated with an independent estimate of early growth rate, female mean size-at-age 3 (*r* =0.44, *R*^2^ = 0.18; Fig. A2). Thus, we determined ω calculated in this manner was a sufficient population-level index of early growth rates.

### Statistical analyses

The magnitudes of SSD among populations based on paired within-lake comparisons of female and male Northern Pike *L*_10_, mean weight, and mean age were evaluated using paired *t*-tests and linear regression to determine deviation from the 1:1 line of no difference. SSD response variables (i.e., SDIs for *L*_10_, mean weight, and mean age) were then tested for relationships with environmental variables (Cisco CPUE, lake surface area, latitude) and life-history traits (ω and *Z*) separately using linear models (LMs) in R v.3.6.2 (R Core Team, 2019). Cisco CPUE and lake surface area were log_10_ transformed to reduce the influence of potential outliers and to normalize distributions. Predictor variables were standardized (Z-scored) by subtracting the mean of all data points from each individual data point and dividing those points by the standard deviation of all data points. With predictor variables standardized to comparable ranges, larger parameter estimates represent greater impacts on response variables for similar changes among predictor variables (Quinn and Keough, 2002). Predictor variables were evaluated for multicollinearity using Pearson’s correlation coefficients (Table A4) and variance inflation factors (VIFs). The VIF values for all covariates were < 3 and thus no predictor variables were initially removed on this basis (Zuur *et al*., 2010). Models of SSD with both environmental and life-history variables were considered independent evaluations of hypotheses, and the evaluation of each response variable (SDIs for *L*_10_, mean weight, and mean age) was adjusted separately for multiple testing with the false discovery rate (FDR) controlling procedure of Benjamini and Hochberg (1995) at a 5% FDR level.

## RESULTS

### Degree of female-biased SSD

Northern Pike mean length-at-age across Ontario confirmed that the body sizes of female and male Northern Pike are similar early in life, while differences in length-at-age were greatest in older age classes (Fig. 2, Table A5). Female-male differences in asymptotic size (*L*_10_:173.02 mm, paired *t*-test, *t*_101_ = 20.82, *P* < 0.0001; Fig. 3A), mean weight (691.08 g, paired *t*-test, *t*_101_ = 13.64, *P* < 0.0001; Fig. 3B), and mean age (0.70 years, paired *t*-test, *t*_101_ = 9.11, *P* < 0.0001; Fig. 3C) were consistently and significantly different, being greater for females than males in each case.

**Fig. 2.**
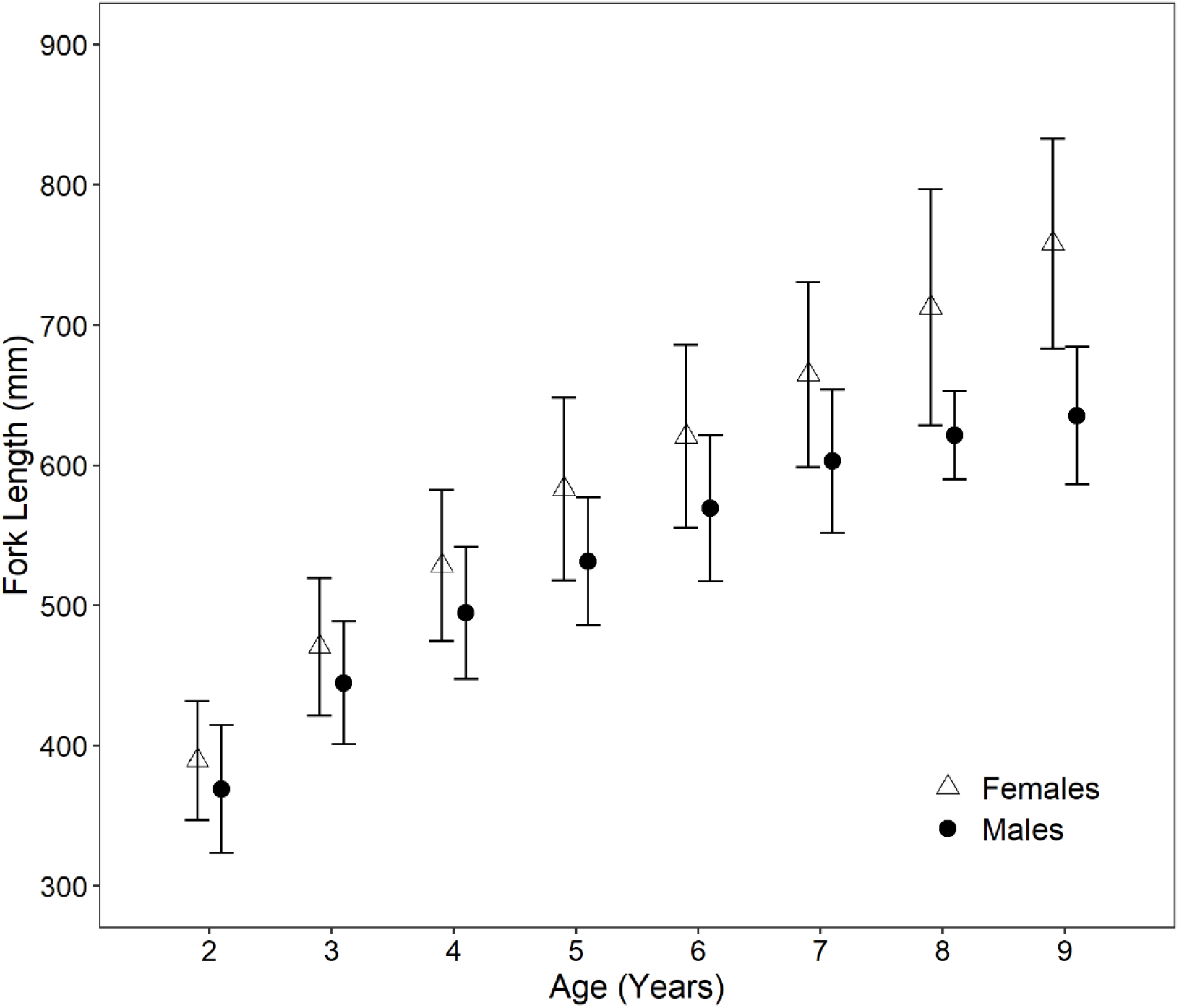
Mean fork length (mm) at age (years) for female and male Northern Pike in the FWIN dataset ± one standard deviation. Means were calculated using means from individual populations (i.e., lakes) with at least three individuals in an age class.

**Fig. 3.**
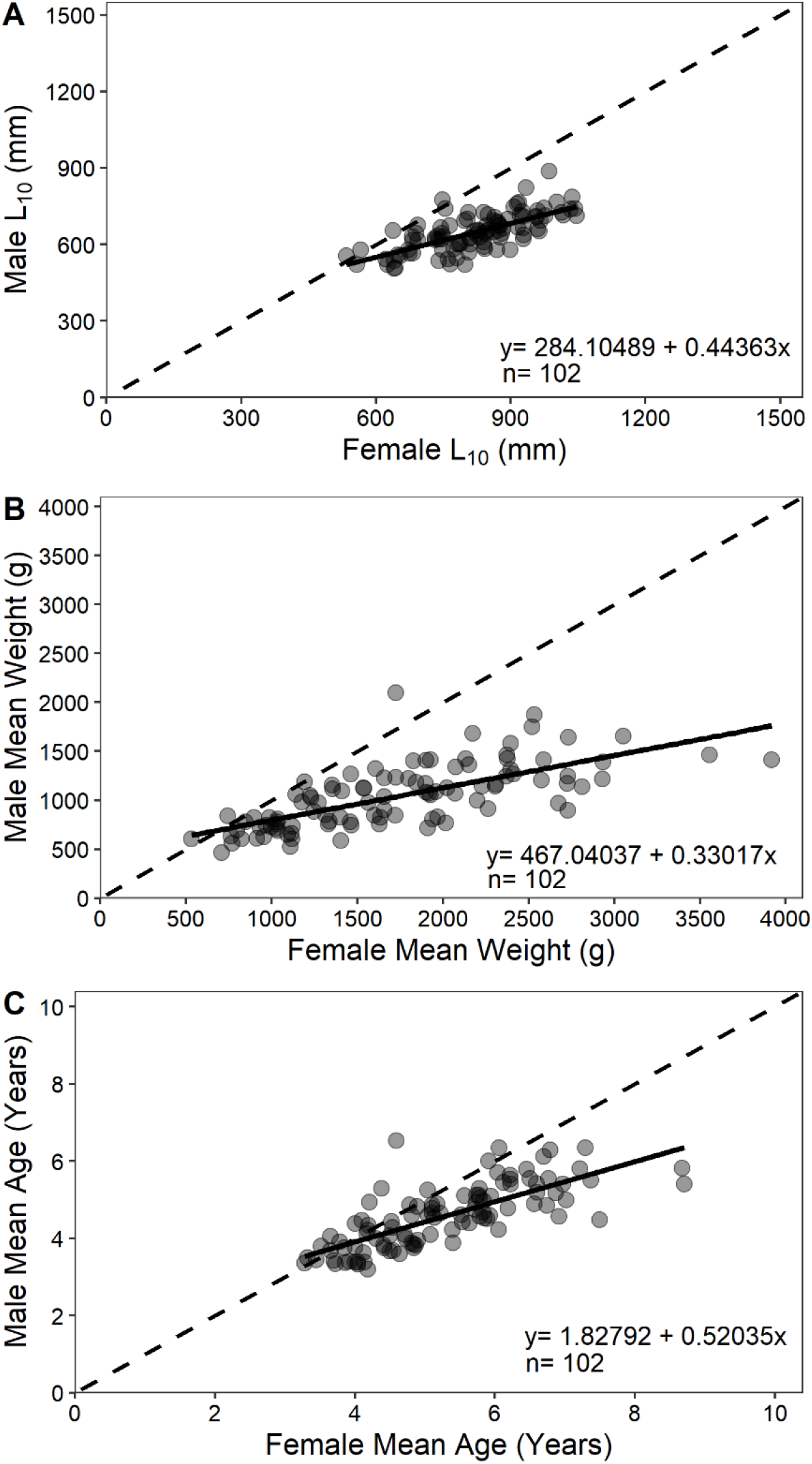
Linear relationships between (A) *L*_10_ (mm), (B) mean weight (g), and (C) mean age (years) of paired (i.e., from the same lake) female and male Northern Pike populations (*F*_100_ = 91.94 to 126.90, *P* < 0.0001). The solid lines represent the linear relationships, and the dashed lines represent 1:1 relationships (i.e., no SSD). The equations for the linear regressions and the sample size (*n*) of lakes are shown.

### Effects of environmental variables on SSD

The degree of female-biased SSD was partially related to environmental variables and depended on the response variable tested. The SDI for *L*_10_ (LM, *df* = 98, *F* = 0.71, *P* = 0.55, *R*^2^ = 0.02) was not related to Cisco abundance, lake surface area, or latitude (*P* > 0.05 for each parameter; Table A6). However, the SDI for mean weight (LM, *df* = 98, *F* = 4.69, *P* = 0.004, *R*^2^ = 0.13) was positively related to Cisco abundance (*P* = 0.02; Fig. 4A).

**Fig. 4.**
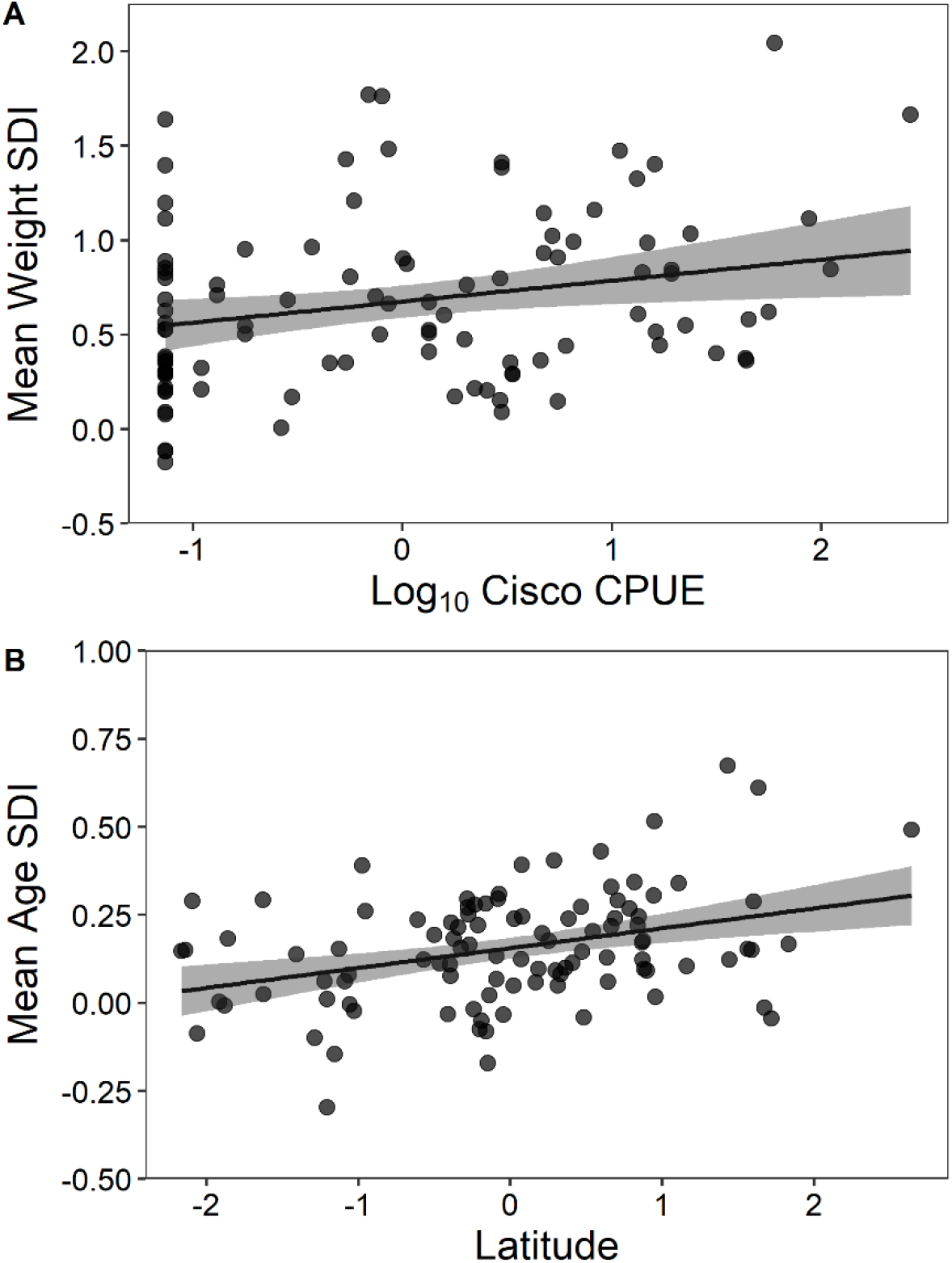
Relationship between (A) mean weight SDI and Cisco CPUE (*P* = 0.02) and (B) mean age SDI and latitude (*P* = 0.001). Cisco CPUE and latitude are standardized (Z-scored). Shading represents 95% confidence intervals around the regression line.

Additionally, the SDI for mean age (LM, *df* = 98, *F* = 5.60, *P* = 0.001, *R*^2^ = 0.15) was positively related to latitude (*P* = 0.001; Fig. 4B). The SDIs for mean weight and mean age were not significantly related to any other environmental variables.

### Life history associations with SSD

The degree of female-biased SSD in Northern Pike was strongly related to life-history traits. The SDI for *L*_10_ (LM, *df* = 98, *F* = 12.49, *P* < 0.0001, *R*^2^ = 0.20) was negatively related to female Northern Pike early growth rates (*P* = 0.0002; Fig. 5A) but was not related to female Northern Pike mortality rates (*P* > 0.05; Table A7). In contrast, the SDI for mean weight (LM, *df* = 98, *F* = 34.76, *P* < 0.0001, *R*^2^ = 0.42) was negatively related to both female Northern Pike mortality and early growth rates (*P* = 0.0002 and < 0.0001, respectively; Fig. 5B,C). Finally, the SDI for mean age (LM, *df* = 98, *F* = 31.91, *P* < 0.0001, *R*^2^ = 0.39) was negatively related to both female Northern Pike mortality and early growth rates (*P* = 0.002 and < 0.0001, respectively; Fig. 5D,E).

**Fig. 5.**
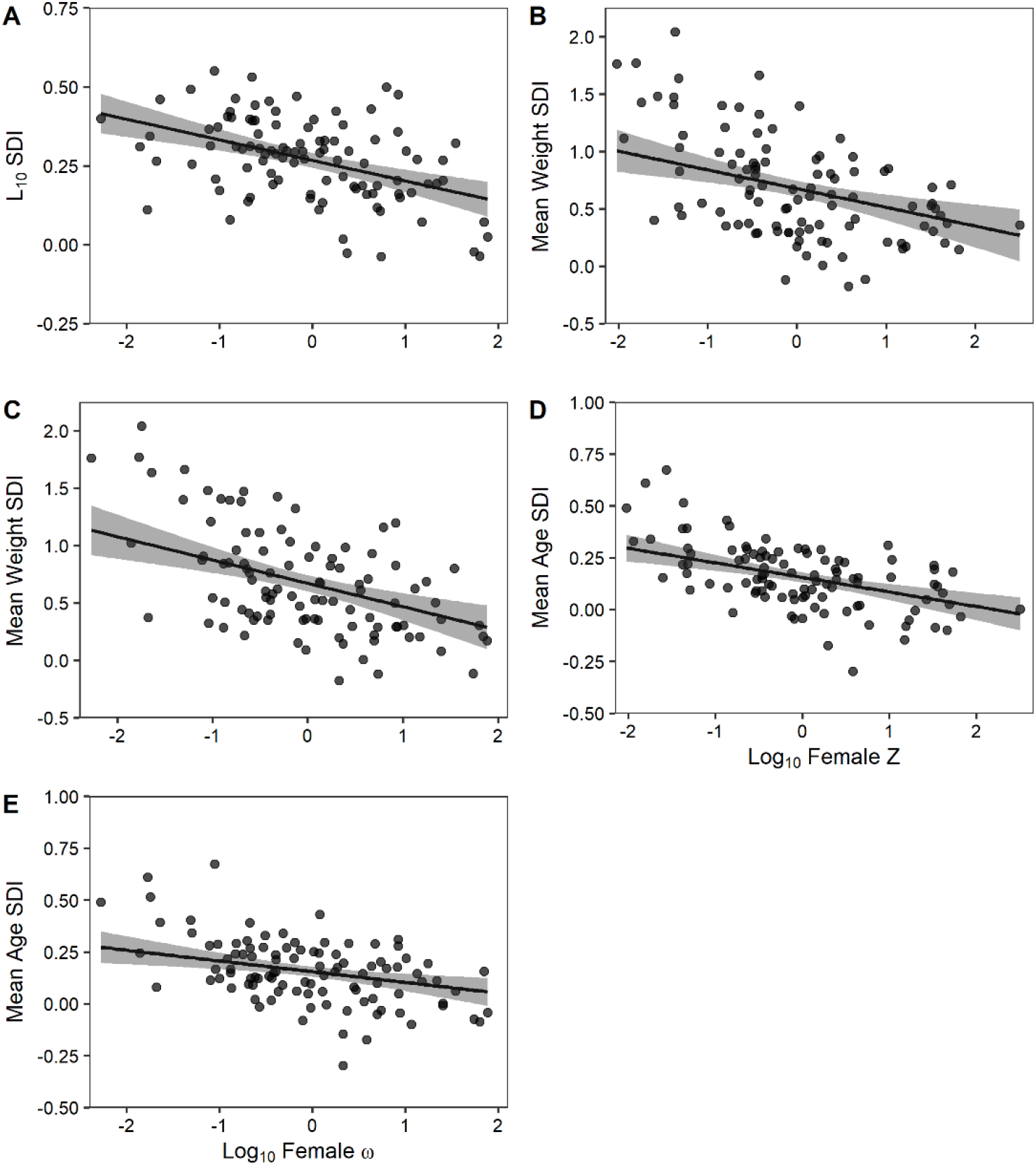
Relationship between (A) *L*_10_ SDI and female Northern Pike ω (*P* = 0.0002), (B) mean weight SDI and female Northern Pike *Z* (*P* = 0.0002), (C) mean weight SDI and female Northern Pike ω (*P* < 0.0001), (D) mean age SDI and female Northern Pike *Z* (*P* < 0.0001), and (E) mean age SDI and female Northern Pike ω (*P* = 0.002). Female Northern Pike ω and *Z* are standardized (Z-scored). Shading represents 95% confidence intervals as in Fig. 4.

## DISCUSSION

The degree of female-biased SSD in Northern Pike increased with age, and variation in SSD among populations was related to both environmental and life-history traits. Our hypotheses that female-biased SSD in Northern Pike (more pronounced differences between females and males) would be greater in higher-latitude lakes with greater Cisco densities were supported in part by a positive relationship between Cisco CPUE and female-biased SSD in mean weight. Additionally, differences between female and male mean age were positively related to latitude, supporting the assertion that lakes at higher latitudes may exhibit longer-living female Northern Pike relative to males. Our hypotheses that Northern Pike populations with lower female mortality and early growth rates would exhibit greater female-biased SSD were also supported; female-biased SSD in mean weight was negatively related to both female Northern Pike mortality and early growth rates, and female-biased SSD in asymptotic size was negatively related to female Northern Pike early growth rates. Further, differences in female and male mean age were negatively related to both female Northern Pike mortality and early growth rates.

Greater female-biased SSD in mean weight appears to result from an increase in high energy prey availability (i.e., Cisco). An increase in surplus energy, particularly at larger sizes, provides additional energy toward both somatic growth and reproduction (e.g., gonadal development), which are reflected more strongly in greater mean weights (which respond more rapidly to changing environmental conditions) than in mean lengths (which are more fixed than weight given the permanence of skeletal vs. soft tissues). Further, larger females with larger gape will be better positioned to take advantage of large-bodied and energy-rich prey, leading to more efficient post-maturation growth relative to smaller males (Pazzia *et al*., 2002; Giacomini *et al*., 2013). Thus, initially small differences in body size between the sexes may quickly become exacerbated with the availability of large-bodied prey. Moreover, large, energy-dense prey availability may promote more efficient physiological maintenance, resulting in longer living populations (Kirkwood and Rosen, 1991).

Previous studies have shown Northern Pike mortality to be greater in lower-latitude Ontario lakes experiencing warmer thermal environments and greater human population density (Griffiths *et al*., 2004; Kennedy *et al*., 2018), which may also help explain greater SSD in mean weight and age at higher latitudes. As water temperature is an important and positive correlate of both growth and natural mortality in fish (Pauly, 1980; Heino *et al*., 2015; Honsey *et al*., 2019), warmer lakes at lower latitudes may support Northern Pike populations with faster life histories (Griffiths *et al*., 2004). This more rapid life history might leave less scope for differences between female and male mean body size and age (i.e., reduced SSD). Such plasticity in SSD has similarly been shown in other poikilothermic species along temperature and latitudinal gradients (Morbey, 2018; Tarr *et al*., 2019). Additionally, lower-latitude lakes closer to urban population centers of southern Ontario likely experience greater fishing pressure and fishing-induced mortalities (Post and Parkinson, 2012). In response to higher mortality, populations at lower latitudes may also reach maturity earlier to maximize lifetime reproductive fitness (Heino *et al*., 2015), reducing the opportunity for SSD by allocating energy to reproduction instead of somatic growth earlier and at smaller sizes than northern populations.

The degree of female-biased SSD was negatively related to both female mortality and early growth indices. Contrary to our observed relationship between mortality and SSD, recent research on female-biased SSD in Lake Whitefish revealed that SSD was *positively* related to the mortality rate of populations [higher mortality associated with greater SSD (Morbey, 2018)]. According to Parker (1992), female-biased SSD should be sensitive to mortality rate, such that low-mortality environments should promote the evolution of later maturation and slower early growth rates of both sexes. This can result in “von Bertalanffy buffering” in some species, where sexes mature later at similar sizes closer to asymptotic size, thus reducing the potential for SSD to manifest (Morbey, 2018). While we were unable to estimate size at maturity in our dataset, mortality is known to relate inversely with size at maturity (Reznick *et al*., 1990). Thus, we may predict based on our findings that female Northern Pike mature later and at larger sizes than males under low mortality conditions, resulting in greater female-biased SSD. This suggests that female-biased SSD in Northern Pike is not limited by von Bertalanffy buffering but is rather due to other processes. This pattern may be further facilitated by male Northern Pike mortality being typically elevated relative to females from the same population (Fig. A3). Additionally, rapid early growth is often associated with smaller adult sizes, younger ages at maturation, and shorter lifespans for many species (Lemaître *et al*., 2015). Thus, more rapid female early growth relative to males may limit the capacity for female-biased SSD in Northern Pike.

Across all populations, female-biased SSD in Northern Pike increased with age, as observed in several other species displaying SSD (e.g., Rennie *et al*., 2008). This strongly suggests energetic (and potentially behavioral) differences between males and females post-maturation. Theoretically, female Northern Pike should experience greater selective pressure to reach larger body sizes than males because their reproductive fitness is more strongly related to body size (Parker, 1992). Reaching larger body sizes increases fecundity in females, and larger numbers of eggs is correlated with better offspring survival (Craig and Kipling, 1983). As such, females may exhibit sex-specific behavior (e.g., foraging strategies) that promote larger body sizes. Female Northern Pike have been found to forage more frequently than males (Diana, 1983), and larger Northern Pike were found to be more mobile and inhabit deeper habitats of lakes relative to smaller Northern Pike (Chapman and Mackay, 1984; Pierce and Tomcko, 2005). Thus, females may forage on large, energy-dense prey (i.e., Cisco) more so than males leading to larger adult body sizes and lower mortality rates (Kennedy *et al*., 2018). Similar feeding behavior has similarly been hypothesized to explain SSD in Walleye (Kaufman *et al*., 2009; Lepak *et al*., 2012).

While foraging behavior promoting larger body size is likely apparent in both sexes, selective pressures on reproductive success should result in females seeking the benefits of foraging more frequently and efficiently on offshore prey (i.e., Cisco) than males. There are risks associated with activity in offshore habitats with less cover from predators, including other cannibalistic Northern Pike (Giles *et al*., 1986). If taking such risks in exposure during foraging does not substantially improve male reproductive success, then selective pressures on males to forage more frequently in offshore habitats should be low. Males might also take such risks if Northern Pike were size-selective breeders (e.g., larger females breed with larger males), had high levels of sperm competition, or if males displayed protective behavior over developing eggs (Roff, 1983; Parker, 1992). However, we have found no evidence in the literature of size-selective breeding or protective behavior in Northern Pike, and Northern Pike have limited sperm competition due to distinct pairing between females and males; female Northern Pike are often found to be accompanied by only one to three males during spawning (Clark, 1950). Thus, escaping vulnerability to predation (via rapid early growth) and reaching maturity should be the primary selection forces on male Northern Pike growth and behavior. This presumed lower level of activity of males has been shown in other fish species exhibiting female-biased SSD (i.e., Walleye and Yellow Perch) via bioenergetic modeling, where males were estimated to have lower consumption rates, metabolic costs, and conversion efficiencies than females (Rennie *et al*., 2008). Additionally, males may experience mating opportunity advantages of earlier maturation, which could result in them maturing at smaller sizes relative to females (Parker, 1992).

As a communal spawning species, we expected female Northern Pike body size to vary more than male body size (Pyron *et al*., 2013). Indeed, female Northern Pike had significantly larger body sizes than males, and this difference increased with age. This highlights the importance of fecundity selection on Northern Pike life histories and populations. As Northern Pike are cannibalistic and commonly inhabit systems with other top predators, they experience high juvenile mortality rates from predation (Kipling and Frost, 1970; Grimm and Klinge, 1996). High juvenile mortality is a strong selection force for large fecund female Northern Pike, and our study makes it increasingly apparent that there are sex-specific selection pressures on growth, maturity, foraging, and habitat selection in this species. Our results suggest that future studies into Northern Pike ecology and evolution should explicitly consider the sex-specific benefits and costs of larger size where possible.

This study on female-biased SSD in Northern Pike revealed significantly less SSD in fast-growing populations with high mortality rates. Higher latitudes were positively related to the difference in female and male mean age, and Cisco density was positively related to female-biased SSD in mean weight, suggesting that colder temperatures and Cisco availability may be key drivers of longevity and post-maturation growth needed for greater female-biased SSD. This is a particularly important finding, as cold-water Cisco populations are currently declining in their southern range due to climate change and eutrophication (Jacobson *et al*., 2012). The loss of key cold-water prey like Cisco may therefore substantially impact the availability of “trophy”-sized Northern Pike and potentially negatively affect recreational fishing opportunities for this species. Furthermore, fisheries managers should continue to incorporate sex differences in body size when setting fishing regulations for Northern Pike (e.g., slot size limits). The results of this study provide novel insight to potential linkages between Northern Pike ecology and life-history outcomes and is the first we know of to explain variability in female-biased SSD among Northern Pike populations by both environmental and life-history variables across a large geographic range.

## ACKNOWLEDGMENTS

We thank George Morgan and Dr. Cindy Chu from the Ontario Ministry of Natural Resources and Forestry for providing data, as well as John Gunn for discussions that helped initiate the project. We also thank Dr. Brian Shuter, anonymous reviewers, and Dr. Derek Roff for their constructive and helpful reviews on earlier versions of the manuscript. This work was supported by grants from the Rainy Lakes Fisheries Charity Trust, NSERC Discovery and the Canada Research Chairs Program to M.D.R., and from the IISD-Experimental Lakes Area to M.D.R. and P.K. There are no conflicts of interest with this manuscript.

## DATA ACCESSIBILTY

All data presented are available on request from the Ontario Ministry of Natural Resources and Forestry.

## APPENDIX

Descriptive statistics and correlation tables for response and predictor variables, example calculation of *S* and *Z* estimates, model outputs for Northern Pike SSD analyses with environmental and life-history variables, and the relationships between female and male *Z* and ω estimates.

**Table A1.**
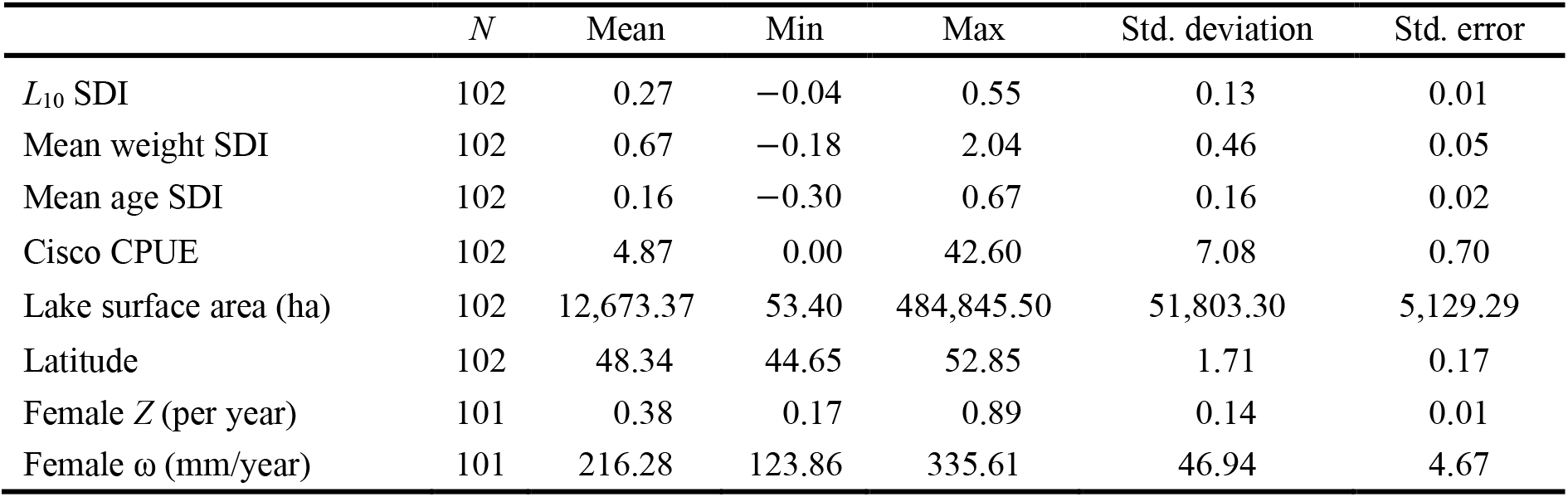
Descriptive statistics for Northern Pike SSD response variables and environmental and life-history predictor variables included in linear models. Negative values for SDI indicate male-biased SSD, whereas positive values indicate female-biased SSD.

**Table A2.** Pearson correlation coefficients among Northern Pike SSD response variables.

| | $L_{10}$ SDI | Mean weight SDI | Mean age SDI |
| --- | --- | --- | --- |
| $L_{10}$ SDI | 1.00 | 0.62 | 0.50 |
| Mean weight SDI | 0.62 | 1.00 | 0.83 |
| Mean age SDI | 0.50 | 0.83 | 1.00 |

**Table A3.**
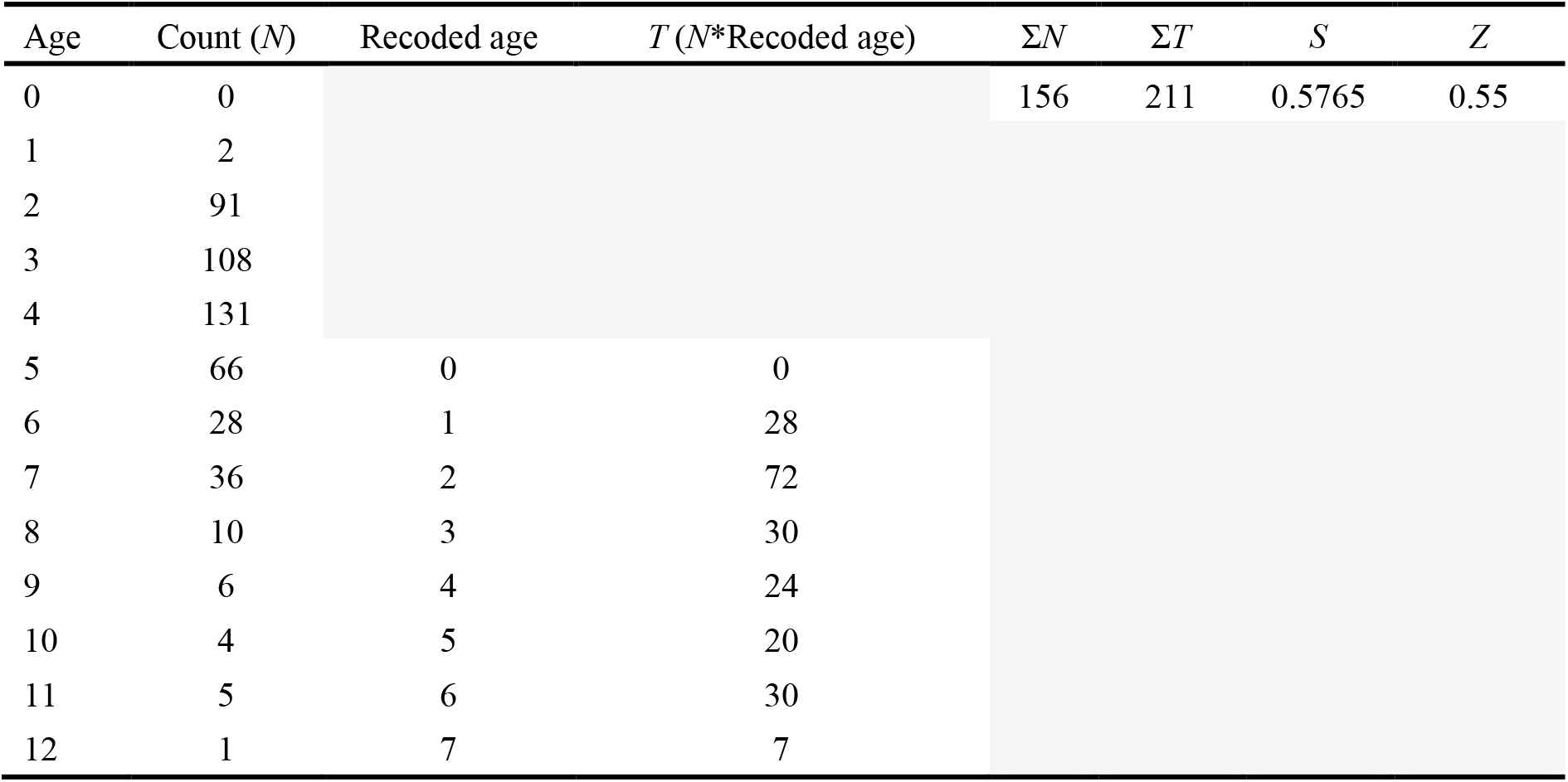
Example calculation of *S* and *T* estimates for female Northern Pike in Lake Nipissing, Ontario. Σ*N* only includes counts for the modal age class (age 5) and older age classes.

| Age | Count ($N$) | Recoded age | $T$ ($N$*Recoded age) | $\Sigma N$ | $\Sigma T$ | $S$ | $Z$ |
| --- | --- | --- | --- | --- | --- | --- | --- |
| 0 | 0 |  |  | 156 | 211 | 0.5765 | 0.55 |
| 1 | 2 |  |  |  |  |  |  |
| 2 | 91 |  |  |  |  |  |  |
| 3 | 108 |  |  |  |  |  |  |
| 4 | 131 |  |  |  |  |  |  |
| 5 | 66 | 0 | 0 |  |  |  |  |
| 6 | 28 | 1 | 28 |  |  |  |  |
| 7 | 36 | 2 | 72 |  |  |  |  |
| 8 | 10 | 3 | 30 |  |  |  |  |
| 9 | 6 | 4 | 24 |  |  |  |  |
| 10 | 4 | 5 | 20 |  |  |  |  |
| 11 | 5 | 6 | 30 |  |  |  |  |
| 12 | 1 | 7 | 7 |  |  |  |  |

**Table A4.** Pearson correlation coefficients among predictor variables included in the analysis.

| | Log<sub>10</sub><br>female Z | Log<sub>10</sub><br>female $\omega$ | Log<sub>10</sub><br>Cisco CPUE | Log<sub>10</sub><br>surface area | Latitude |
| --- | --- | --- | --- | --- | --- |
| Log<sub>10</sub> female Z | 1.00 | 0.56 | -0.48 | -0.28 | -0.55 |
| Log<sub>10</sub> female $\omega$ | 0.56 | 1.00 | -0.38 | -0.24 | -0.47 |
| Log<sub>10</sub> Cisco CPUE | -0.48 | -0.38 | 1.00 | 0.37 | 0.32 |
| Log<sub>10</sub> surface area | -0.28 | -0.24 | 0.37 | 1.00 | 0.33 |
| Latitude | -0.55 | -0.47 | 0.32 | 0.33 | 1.00 |

**Table A5.** Summary statistics for the mean fork length (mm) at age for populations with at least three individuals in each age class. *N* represents the number of lakes the estimate is derived from.

| Sex | Age | *N* | Fork length (mm) | Std. error | Std. deviation | CI |
| --- | --- | --- | --- | --- | --- | --- |
| Female | 2 | 94 | 389.25 | 4.36 | 42.28 | 8.66 |
|  | 3 | 217 | 470.46 | 3.32 | 48.93 | 6.55 |
|  | 4 | 224 | 528.22 | 3.60 | 53.87 | 7.09 |
|  | 5 | 198 | 582.93 | 4.63 | 65.16 | 9.13 |
|  | 6 | 130 | 620.57 | 5.71 | 65.10 | 11.30 |
|  | 7 | 105 | 664.60 | 6.45 | 66.09 | 12.79 |
|  | 8 | 71 | 712.51 | 10.01 | 84.37 | 19.97 |
|  | 9 | 34 | 758.06 | 12.82 | 74.77 | 26.09 |
| Male | 2 | 94 | 368.85 | 4.69 | 45.50 | 9.32 |
|  | 3 | 200 | 444.74 | 3.09 | 43.70 | 6.09 |
|  | 4 | 222 | 494.87 | 3.17 | 47.23 | 6.25 |
|  | 5 | 171 | 531.35 | 3.48 | 45.51 | 6.87 |
|  | 6 | 118 | 569.14 | 4.80 | 52.18 | 9.51 |
|  | 7 | 60 | 602.89 | 6.61 | 51.24 | 13.24 |
|  | 8 | 25 | 621.32 | 6.27 | 31.35 | 12.94 |
|  | 9 | 7 | 635.39 | 18.54 | 49.05 | 45.36 |

**Table A6.** Linear model outputs for the effects of environmental predictor variables on Northern Pike SDI for *L*_10_ (*df* = 98, *F* = 0.71, *P* = 0.55, *R*^2^ = 0.02), mean weight (*df* = 98, *F* = 4.69, *P* = 0.004, *R*^2^ = 0.13), and mean age (*df* = 98, *F* = 5.60, *P* = 0.001, *R*^2^ = 0.15). Significant predictor variables are displayed in **bold**. Benjamini-Hochberg critical values with an FDR of 5% [(l/m)Q (5%)] are provided. Predictor variables have been standardized (Z-scored).

| Response variable | Predictor variable(s) | df | Parameter estimate | Std. error | t | P | (1/m)Q (5%) | R<sup>2</sup> |
| --- | --- | --- | --- | --- | --- | --- | --- | --- |
| $L_{10}$ SDI | Log<sub>10</sub> Cisco CPUE | 98 | 0.006 | 0.015 | 0.394 | 0.694 | | |
|  | Log<sub>10</sub> lake surface area | 98 | 0.011 | 0.015 | 0.728 | 0.468 |  |  |
|  | Latitude | 98 | 0.009 | 0.014 | 0.628 | 0.532 |  |  |
|  |  |  |  |  |  | **0.055** | **0.050** | **0.021** |
| Mean weight SDI | **Log<sub>10</sub> Cisco CPUE** | **98** | **0.110** | **0.048** | **2.281** | **0.025** |  |  |
|  | Log<sub>10</sub> lake surface area | 98 | 0.004 | 0.048 | 0.092 | 0.927 |  |  |
|  | Latitude | 98 | 0.088 | 0.047 | 1.857 | 0.066 |  |  |
|  |  |  |  |  |  | **0.004** | **0.033** | **0.126** |
| Mean age SDI | Log<sub>10</sub> Cisco CPUE | 98 | 0.024 | 0.016 | 1.485 | 0.141 |  |  |
|  | Log<sub>10</sub> lake surface area | 98 | -0.015 | 0.017 | -0.917 | 0.361 |  |  |
|  | **Latitude** | **98** | **0.054** | **0.016** | **3.300** | **0.001** |  |  |
|  |  |  |  |  |  | **0.001** | **0.017** | **0.146** |

**Table A7.** Linear model outputs for the effects of female Northern Pike *Z* and ω on Northern Pike SDI for *L*_10_ (*df* = 98, *F* = 12.49, *P* = 1.48e-05, *R*^2^ = 0.20), mean weight (*df* = 98, *F* = 34.76, *P* = 3.89e-12, *R*^2^ = 0.42), and mean age (*df* = 98, *F* = 31.91, *P* = 2.13e-11, *R*^2^ = 0.39). Significant predictor variables are displayed in **bold**. Benjamini-Hochberg critical values with an FDR of 5% [(l/m)Q (5%)] are provided. Predictor variables have been standardized (Z-scored).

| Response variable | Predictor variable(s) | df | Parameter estimate | Std. error | t | P | (1/m)Q (5%) | R<sup>2</sup> |
| --- | --- | --- | --- | --- | --- | --- | --- | --- |
| $L_{10}$ SDI | Log<sub>10</sub> female Z | 98 | -0.010 | 0.014 | -0.687 | 0.494 | | |
|  | **Log<sub>10</sub> female $\omega$** | **98** | **-0.060** | **0.015** | **-3.891** | **1.82e-04** |  |  |
|  |  |  |  |  |  | **1.48e-05** | **0.050** | **0.203** |
| Mean weight SDI | **Log<sub>10</sub> female Z** | **98** | **-0.163** | **0.043** | **-3.809** | **2.43e-04** |  |  |
|  | **Log<sub>10</sub> female $\omega$** | **98** | **-0.203** | **0.046** | **-4.400** | **2.76e-05** |  |  |
|  |  |  |  |  |  | **3.89e-12** | **0.017** | **0.415** |
| Mean age SDI | **Log<sub>10</sub> female Z** | **98** | **-0.070** | **0.015** | **-4.656** | **1.01e-05** |  |  |
|  | **Log<sub>10</sub> female $\omega$** | **98** | **-0.052** | **0.016** | **-3.170** | **0.002** |  |  |
|  |  |  |  |  |  | **2.13e-11** | **0.033** | **0.394** |

**Fig. A1.**
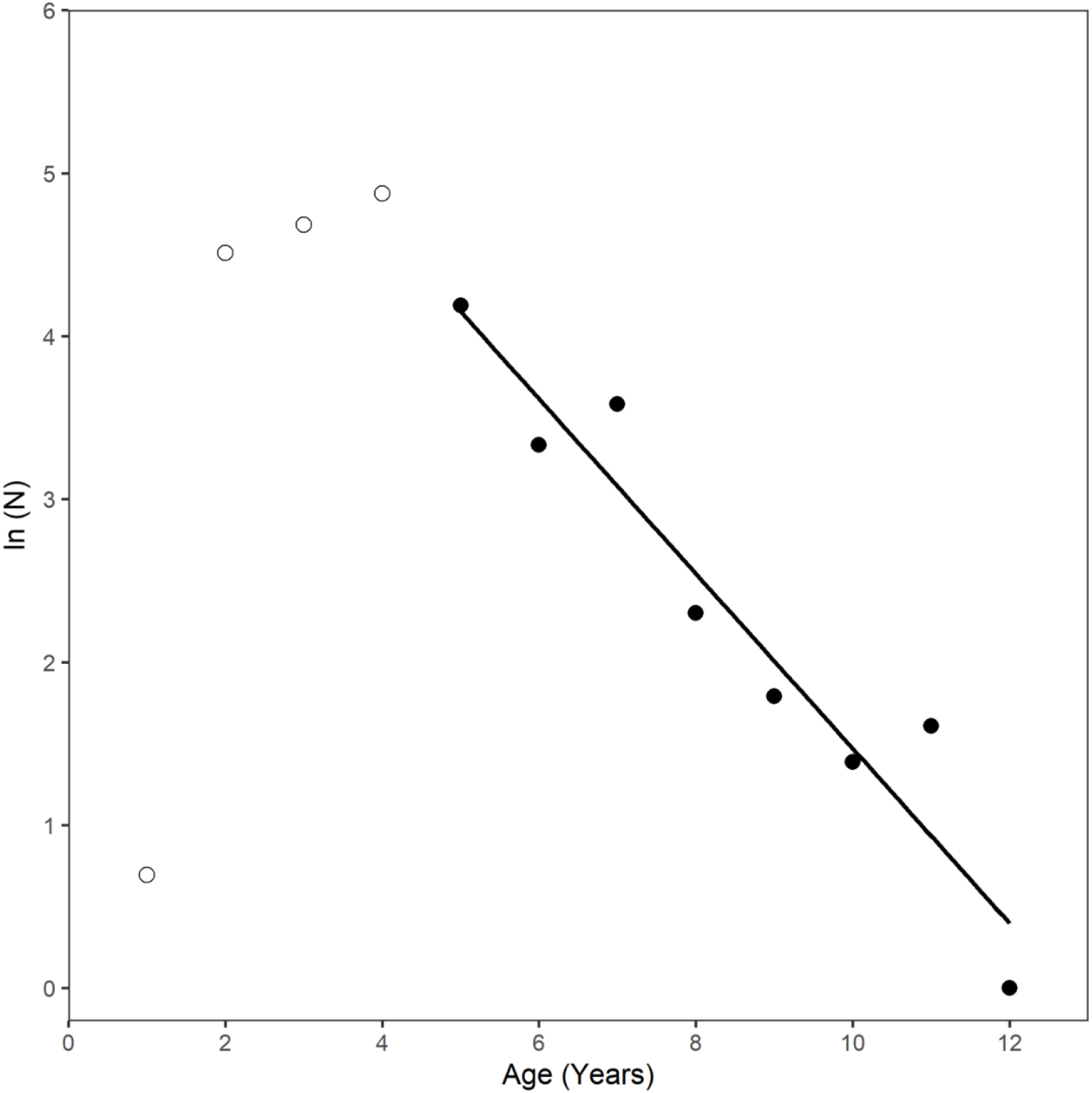
Example of the descending limb of a catch curve used to estimate instantaneous total mortality for female Northern Pike sampled in Lake Nipissing, Ontario. Solid dots represent the log number of individuals caught in each age class on the descending limb of the catch curve. The solid line is a regression line.

**Fig. A2.**
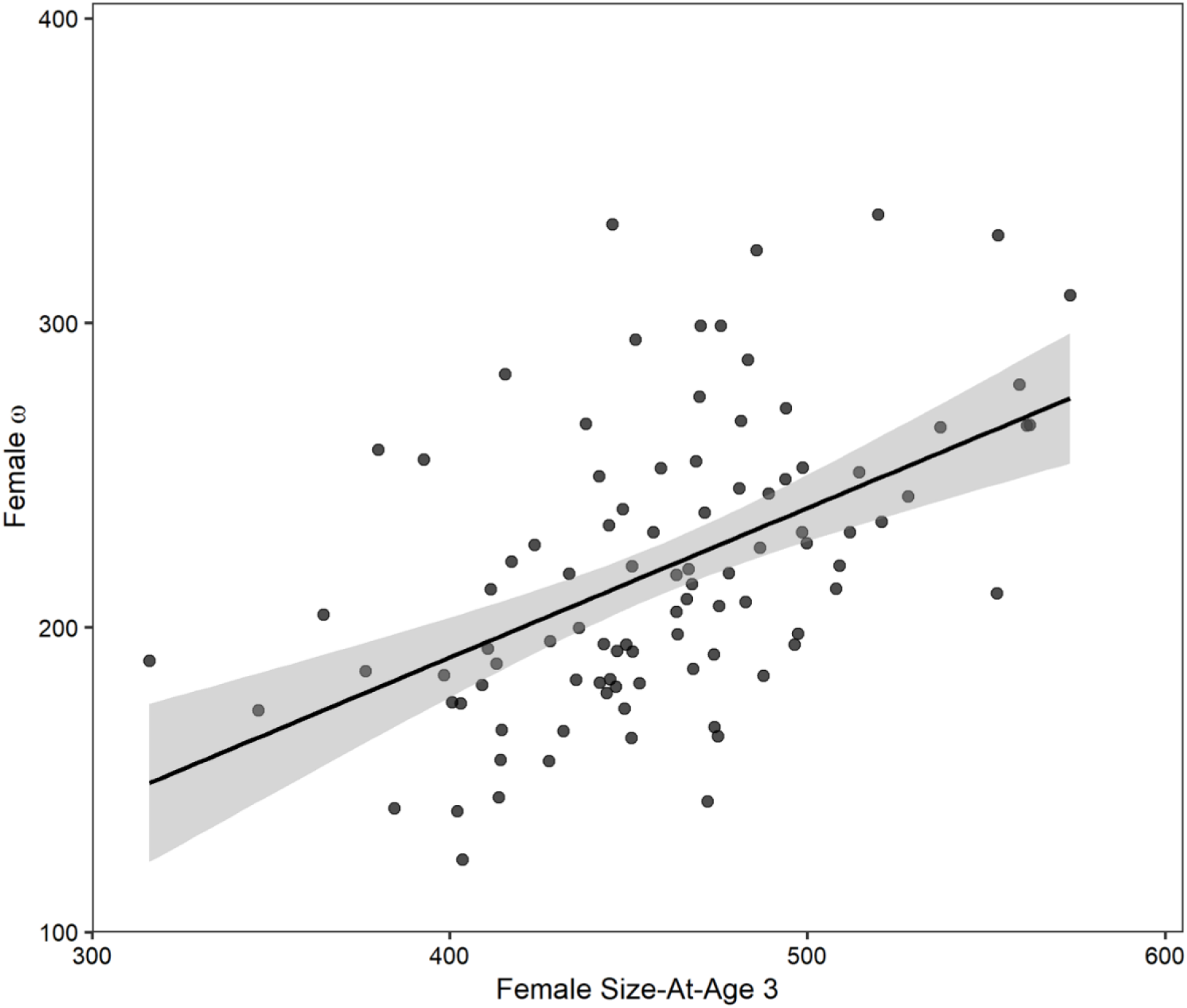
Relationship between female Northern Pike early growth rate estimates (ω) and size-at-age 3 (mm) (LM, *y* = 0.57*x* + 38.11, *df* = 94, *t* = 4.73, *P* < 0.0001, *R*^2^ = 0.18). Shading represents 95% confidence intervals around the regression line.

**Fig. A3.**
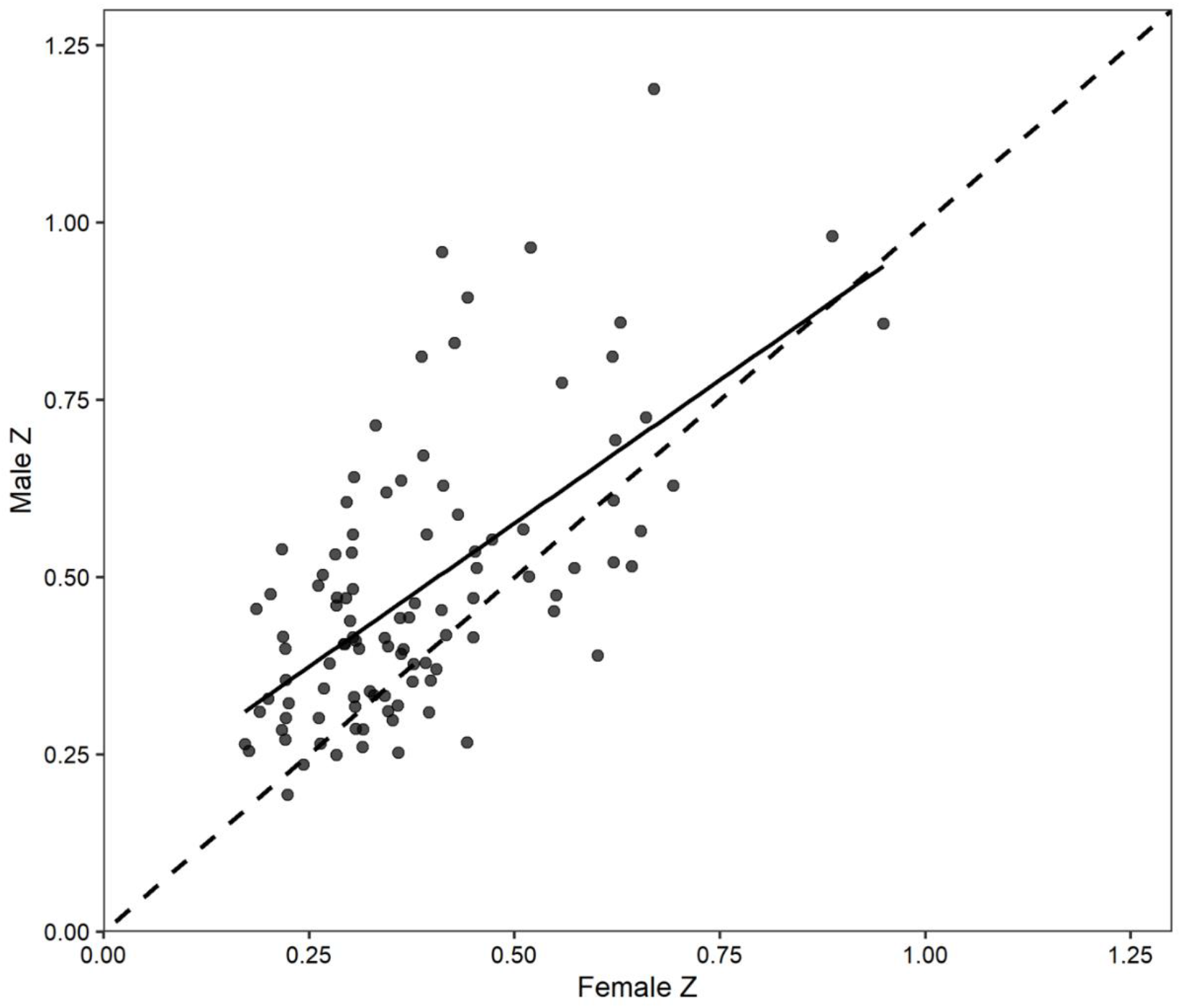
Relationship between male and female Northern Pike instantaneous total mortality estimates (*Z*; LM, *y* = 0.81*x* + 0.17, *df* = 100, *t* = 8.12, *P* < 0.0001, *R*^2^ = 0.39). The dashed line indicates a 1:1 relationship.

